# Bacteriology and Antibiotic Prescription Patterns in a Malawian Tertiary Hospital Burns Unit

**DOI:** 10.1101/419713

**Authors:** Stephen Kasenda, Donnie Mategula, Geoffrey Elihu Manda, Tilinde Keith Chokotho

**Author notes:** **Competing interests** The authors declare that they have no competing interests.

## Abstract

**Introduction:** Infections are responsible for up to 85% of deaths in patients with burn injuries. Proper management of infections in patients with burns requires knowledge of local microbial landscape and antimicrobial resistance patterns. Most burns units in low to middle income countries lack this data to guide patient management.

**Methods and results:** We conducted a retrospective audit of adult (≥17 years) patient records admitted between at 1^st^ June 2007 and May 2017 at Queen Elizabeth Central Hospital Burns unit in Blantyre Malawi with an index complaint of burn injury. Descriptive statistical analysis was performed to determine antibiotic prescription patterns, microbial isolates and antimicrobial resistance patterns on the 500 patient files that met the inclusion criteria. Cephalosporin’s and Penicillin’s constituted 72.3% of the 328 antibiotic prescriptions given to 212 patients and 84% of all prescriptions were potentially inappropriate. A total of 102 bacterial isolates were identified and a majority (30.4%; n=31) were resistant to Aminoglycosides and Aminocyclitols (23.5%; n=24); seconded by Penicillin’s at 19.6% (n=20). Pseudomonas, staphylococcus and streptococcus species constituted 36.1%, 25% and 16.7% of all resistant bacteria that were isolated and they were thus the most common bacterial isolates. Drug resistance was more common among gram negative bacteria (48.8% versus 26.2%) and a greater proportion of patients (74.1%) that had antibiotic sensitivity testing were affected by drug resistant gram negative bacteria which appear on the World Health Organisation list of priority pathogens.

**Conclusion:** The results of our preliminary study point towards nosocomial gram negative bacteria which appear on the World Health Organisations list of priority pathogens as the more common sources of antibiotic resistance. This scenario is potentially driven by inappropriate antibiotic prescriptions as well as clinical and laboratory diagnostic imprecision in addition to the universally recognised post burn pathophysiological changes of hypermetabolism and immunosuppression. Improvements in the areas of antimicrobial stewardship, diagnostic capacity and burns related research are needed in order to achieve optimal therapeutic outcomes and resource utilisation.

## INTRODUCTION

Antimicrobial resistance (AMR) is the ability of a microorganism to stop an antimicrobial from working against it. ^1^ There are concerns of increasing AMR worldwide and its reported inverse relationship with national wealth.^2,3^ In low and middle income countries (LMIC’s) where the incidence of infection is highest, the treatment options are limited.^3–7^ In addition to the innate ability of micro-organisms to mutate, the major contributing factors to increasing AMR can be categorised as legislative factors, patient factors as well as diagnosis and prescription related factors. In Sub-Saharan Africa there is deficiency in enforcement of laws to control production quality of antibiotics and dispensing of antimicrobials based on self prescription and for agricultural use.^8^ Hospital laboratories also tend to have limited capacity to reliably conduct antimicrobial sensitivity tests and there are questions surrounding patients’ adherence to prescribed therapy especially after they start feeling better.^3–5,9–11^ Despite the existence of these pertinent issues, there is a dearth of information on AMR in Sub Saharan Africa.^2,12^

Infections are the most important cause of death in burn victims after resolution of the initial shock phase (in the first 24 hours) accounting for 50-84% of all burns related deaths.^13–16^ Burn patients are at increased risk of infection and sepsis due to loss of the skin barrier, reduced production of antimicrobial peptides (Defensins and Cathelicidins) by epithelial keratinocytes due to increased expression of interleukin 10 (IL-10) and CC-chemokine ligand 2 (CCL2) by altered neutrophils (PMN II).^17–19^ Burn victims also have altered adaptive immunity due to suppressed antigen presenting cell (langerhans and dendritic cells) function and vascular thrombosis.^20^ The worldwide trend of increasing AMR amidst competing infections and a limited range of antimicrobials renders AMR surveillance a necessity especially among special groups such as burn victims. We reviewed ward records covering a 10 year period in order to identify the antibiotic prescription patterns as well as AMR prevalence amongst adult patients admitted in the Queen Elizabeth Central Hospital burns Unit. The latest of two studies that were done on this topic in our setting was done over a decade ago and AMR patterns have not been studied since 2001.^21,22^

## METHODS

### Study Design and Setting

The study was a cross sectional audit of records at Queen Elizabeth Central Hospital (QECH) which is the largest referral hospital in Malawi with a bed capacity of over 1000 located in southern region of the country. It is also the location for the first burns unit in the country with a bed capacity of 32.

### Study population

All patients aged 17 years or more with recorded blood culture or wound swab results, having any type of burn injury and admitted in the QECH Burns Unit between 1^st^ June 2007 and May 2017 met the inclusion criteria of this audit. Patients excluded from the study were those who; had no documentation beyond the admission process, were pronounced dead on arrival, presented for the first time with complications of burns, and those with non-burn-related issues.

### Data collection

We searched the surgery department electronic records as well as the ward registers to identify burn patients who met the inclusion criteria. The following type of data was extracted from patient files; admission date, discharge date, demographic data (age, gender, referring facility or residence). Time to hospital presentation, length of hospital stay, antimicrobials prescribed, microbiology tests done (Blood culture, wound swab, malaria parasite microscopy and rapid test results). Where microbiology tests were performed we also recorded the antibiotic sensitivity results.

### Data entry and analysis

Data entry and initial cleaning was done using Microsoft Excel 2007 and the final data cleaning and analysis was done using SPSS 21. We ran descriptive statistics tests on the demographic data. Data on antibiotics received, bacterial isolates and resistance to antimicrobials were categorised as multiple response variables according to the most common items noted on each list. Frequency of each item was tested following this categorisation.

### Ethical approval

Ethical approval was obtained from the College of Medicine Research and Ethics Committee (COMREC); COMREC reference number P.09/17/2275.

## RESULTS

Five hundred patient files met the inclusion criteria, and all were available for retrieval. The patient median age was 32 years (IQR: 25-45 years; range 17-92 years) and 260(52%) of these were male. The median time to hospital presentation was 1 day (IQR; 0.7– 7 days) and the median length of hospital stay was 20 days (IQR; 8 – 47 days; range 7 – 228 days). The median total burn surface area was 12% (IQR: 6.0 – 22.0%; range 0 – 100%). One hundred and ninety-four (37.7%) patients were surgically managed.

A subset of 72 patient files contained results of microbiological tests (wound swab and blood culture only). These patients had a median age of 28.5 years (IQR: 22 - 41 years; range 17 - 88 years) and 41(56.9%) of these were male. The median time to hospital presentation was 4 days (IQR; 1 – 14 days) and the median length of hospital stay was 56 days (IQR; 39.25– 90 days; range 7 – 228 days). The median total burn surface area was 12% (IQR: 6.5 – 22.5%; range 1.0 – 45.3%).

### Microbial landscape

Determination of infection or wound colonisation was done almost exclusively by culture of wound swab samples. The only documented blood culture result that we identified had a record of Staphylococcus aureas but drug sensitivity testing was not done on the sample. Seventy two patients had a wound swab sample collected during the course of their admission.

It was evident from the patient files that wounds swabs were almost always collected when a wound infection was suspected based on new onset fever or wound pus. The earliest time to wound swab collection was 3 days (median 28 days; range: 3 – 85 days) from the date of admission. One hundred and two bacterial isolates were identified from wounds swabs of 61 patients that were collected. Thirty four (55.7%) patients’ wound swab samples had mono-bacterial isolates, 11(15.3%) were sterile swabs and there were no fungal isolates. Wound biopsy for quantitative analysis of microbial load was not done on any of the patients. Methicillin Resistant Staphylococcus aureus was isolated in one instance and it was reported to be sensitive to Chloramphenicol only.

Gram positive bacteria were the most prevalent organisms (59.8%) with Staphylococcus alone affecting 55.7% of patients and accounting for the same percentage of all gram-positive bacteria. Enterobacteriaceae (Proteus species, colliforms and salmonella typhimurium) accounted for 20.6% of all isolates and were thus the second most common organisms and the most prevalent gram negative organisms. Proteus species was the most prevalent Enterobacteriaceae cultured (n=15; 71.4%) and Pseudomonas species were the most prevalent gram negative bacteria cultured (n=14) (Table 1). The identity of 12 (11.8%) bacterial isolates obtained from 12 patients was only stated according to appearance after gram staining and not taxonomic classification.

**Table 1:** Prevalence of resistance to each antibiotic

|  | Bacterial |  | Percent of | % Gram |  |
| --- | --- | --- | --- | --- | --- |
|  |  |  |  | positives | % Gram |
|  | Isolates |  | patients | (N=61) | negatives (N=41) |
|  | N | Percent |  |  |  |
| BACILLUS | 10 | 9.8 | 16.4 | 16.4 |  |
| STREPTOCOCCUS SP. | 11 | 10.8 | 18.0 | 18.0 |  |
| STAPHYLOCOCCUS SP. | 34 | 33.3 | 55.7 | 55.7 |  |
| ENTEROBACTERIACEAE | 21 | 20.6 | 34.4 |  | 51.2 |
| PSEUDOMONAS SP. | 14 | 13.7 | 23.0 |  | 34.1 |
| OTHER BACTERIA | 12 | 11.8 | 19.7 | 9.8 | 14.6 |
| **TOTAL** | **102** | **100** | **167.2** | **100** | **100** |

### Prescription patterns

A total of 328 antibiotic prescriptions were given to 212 patients during their admission in the burns unit. Cephalosporin’s and Penicillin’s constituted 72.3% of all prescriptions whilst the rest of the antibiotics prescribed contributed less than 10% each to the census of prescribed antibiotics, hydrophilic antibiotics (Cephalosporin’s, Penicillin’s and Aminoglycosides) accounted for 79.6% of all prescriptions (Table 2). Antibiotic monotherapy was used for 131 (61.8%) of the patients whilst another 53(25%) were prescribed two antibiotics; 20(9.4%) LJLJ three antibiotics and 8 (3.8%) LJ four antibiotics. Wound swabs were not done for 167 (78.8%) of patients that received antibiotics and six of these had malaria diagnosed by blood film microscopy for which they were also treated with Lumefantrine Artemether. None of the patients that received antibiotics without supporting microbiology results had written clinical justification for their antibiotic prescriptions.

**Table 2:** Antibiotic prescription pattern

|  | Prescriptions |  | Percent of patients |
| --- | --- | --- | --- |
|  | N | Percent |  |
| PENICILLINS | 117 | 35.7% | 55.2% |
| CEPHALOSPORINS | 120 | 36.6% | 56.6% |
| MACROLIDES | 2 | 0.6% | 0.9% |
| FLOUROQUINOLONES | 15 | 4.6% | 7.1% |
| AMINOGLYCOSIDES | 24 | 7.3% | 11.3% |
| NITROIMIDAZOLES | 26 | 7.9% | 12.3% |
| CHLORAMPHENICOL | 14 | 4.3% | 6.6% |
| OTHER | 10 | 3.0% | 4.7% |
| **TOTAL** | **328** | **100.0%** | **154.7%** |

Among those that had a wound swab (n=72), 36(50%) patients received 55 antibiotic prescriptions and the most commonly prescribed classes of antibiotics were Cephalosporin’s (43.6%), Aminoglycosides (25.5%) and Penicillin’s (16.4%). (Table 3) These three antibiotics accounted for 85.5% of all the prescribed drugs and they were prescribed to 66.7%, 38.9% and 25% of patients with wound swab results respectively. The rest of the antibiotics were each prescribed to less than 10% of the same cohort of patients.

**Table 3:** Antibiotic prescription pattern for patients that had a wound swab

|  | Responses |  | Percent of Cases |
| --- | --- | --- | --- |
|  | N | Percent |  |
| PENICILLINS | 9 | 16.4% | 25.0% |
| CEPHALOSPORINS | 24 | 43.6% | 66.7% |
| MACROLIDES | 1 | 1.8% | 2.8% |
| FLOUROQUINOLONES | 3 | 5.5% | 8.3% |
| TETRACYCLINES | 1 | 1.8% | 2.8% |
| AMINOGLYCOSIDES | 14 | 25.5% | 38.9% |
| OTHER ANTIBIOTICS | 3 | 5.5% | 8.3% |
| **TOTAL** | **55** | **100.0%** | **152.8%** |

Among the 11 patients that had sterile wound swabs, six had antibiotic prescriptions whereas two did not receive antibiotics and treatment charts of four patients were missing thus their management could not be determined. The antibiotics prescribed to those with sterile cultures were Cephalosporin (5), Penicillin’s (4) and Macrolides (1).

### Resistance patterns

Thirty six drug resistant isolates were identified from the 27 patients that had a sensitivity test in addition to initial culture of the wound swab sample. Pseudomonas, staphylococcus and streptococcus species were the most prevalent antibiotic resistant species isolated and they constituted 36.1%, 25% and 16.7% of all resistant bacteria respectively. (Table 4) The rest of the isolated bacterial species contributed less than 10% each to the total microbial load. Drug resistant gram negative bacterial isolates were more prevalent (n=20; 55.6%), more commonly associated with drug resistance (48.8% versus 26.2%) and affected greatest number of patients (74.1% versus 59.2%). A majority (n=20; 55.6%) of these samples had one drug resistant bacterial isolate with Pseudomonas species contributing 10 (27.8%) of the resistant mono-isolate samples. This was followed by Enterobacteriaceae (4), Staphylococcus species (4) and Streptococcus species (2)

**Table 4:** Prevalence of resistance

|  | Responses |  | Percent of Cases |
| --- | --- | --- | --- |
|  | N | Percent |  |
| **STAPHYLOCOCCUS** | 9 | 25.0% | 33.3% |
| **STREPTOCOCCUS** | 6 | 16.7% | 22.2% |
| **PROTEUS** | 3 | 8.3% | 11.1% |
| **PSEUDOMONAS** | 13 | 36.1% | 48.1% |
| **COLLIFORMS** | 3 | 8.3% | 11.1% |
| **OTHER** | 2 | 5.6% | 7.4% |
| **TOTAL** | 36 | 100.0% | 133.3% |
| Gram positive bacteria | 16 | 26.2% | 59.2% |
| Gram negative bacteria | 20 | 48.8% | 74.1% |
| **TOTAL** |  |  | 133.3% |

Overall prevalence of drug resistance among the organisms and antibiotics tested ranged from zero percent to 70%. The antibiotics associated with the greatest prevalence of resistance were Aminoglycosides and Aminocyclitols (30.4%), Penicillin’s (20.6%) and Cephalosporin’s (19.6%). Antimicrobials least associated with the resistance were Macrolides (8.8%), Carbapenems (5.9%) and Tetracyclines (1.0%) (Tables 5 & 6). Proportions of resistant bacteria to each prescribed antibiotic suggest an association between frequencies of prescription. It was however not possible to establish the statistical significance of this association due to data sparsity.

**Table 5:**
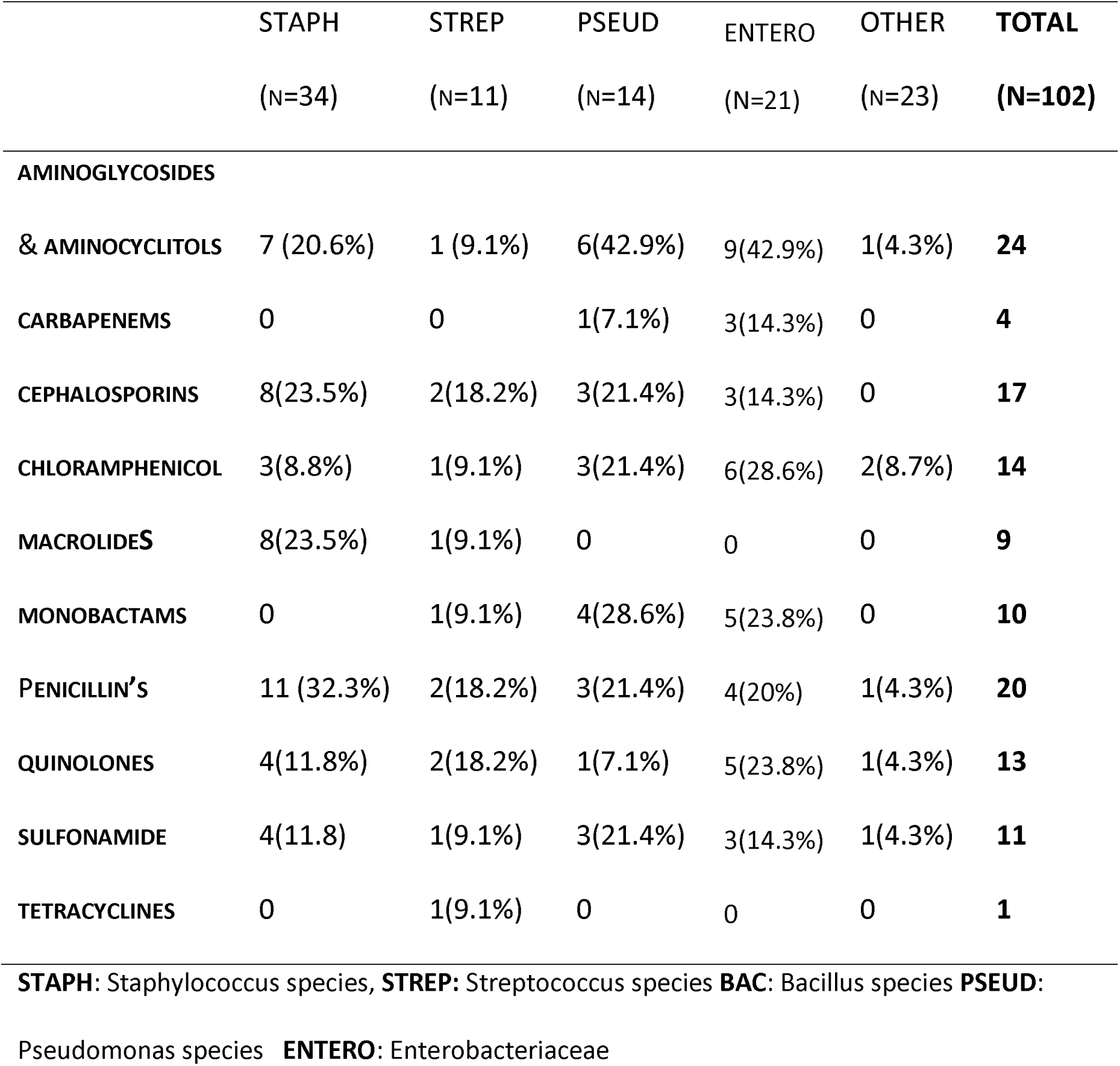
Prevalence of resistance for bacterial isolates and the most commonly used antibiotics

|  | STAPH<br>(N=34) | STREP<br>(N=11) | PSEUD<br>(N=14) | ENTERO<br>(N=21) | OTHER<br>(N=23) | TOTAL<br>(N=102) |
| --- | --- | --- | --- | --- | --- | --- |
| **AMINOGLYCOSIDES** |  |  |  |  |  |  |
| **& AMINOCYCLITOLS** | 7 (20.6%) | 1 (9.1%) | 6(42.9%) | 9(42.9%) | 1(4.3%) | **24** |
| **CARBAPENEMS** | 0 | 0 | 1(7.1%) | 3(14.3%) | 0 | **4** |
| **CEPHALOSPORINS** | 8(23.5%) | 2(18.2%) | 3(21.4%) | 3(14.3%) | 0 | **17** |
| **CHLORAMPHENICOL** | 3(8.8%) | 1(9.1%) | 3(21.4%) | 6(28.6%) | 2(8.7%) | **14** |
| **MACROLIDES** | 8(23.5%) | 1(9.1%) | 0 | 0 | 0 | **9** |
| **MONOBACTAMS** | 0 | 1(9.1%) | 4(28.6%) | 5(23.8%) | 0 | **10** |
| **PENICILLIN'S** | 11 (32.3%) | 2(18.2%) | 3(21.4%) | 4(20%) | 1(4.3%) | **20** |
| **QUINOLONES** | 4(11.8%) | 2(18.2%) | 1(7.1%) | 5(23.8%) | 1(4.3%) | **13** |
| **SULFONAMIDE** | 4(11.8) | 1(9.1%) | 3(21.4%) | 3(14.3%) | 1(4.3%) | **11** |
| **TETRACYCLINES** | 0 | 1(9.1%) | 0 | 0 | 0 | **1** |

**STAPH:** Staphylococcus species, **STREP:** Streptococcus species **BAC:** Bacillus species **PSEUD:**
Pseudomonas species **ENTERO:** Enterobacteriaceae

## DISCUSSION

We conducted a study to determine the prevalent bacterial infections, prescription patterns and AMR patterns among adult patients admitted in the QECH burns unit between 1^st^ June 2007 and 31^st^ May 2017. It was determined that a total of 178(84%) out of the 212 patients that received antibiotics potentially had inappropriate antibiotic prescriptions since they had either sterile wound swab results or no documented bacterial isolates on wound swab or blood culture. Cephalosporin’s and Penicillin’s constituted 72.3% of all prescriptions with each one of the two antibiotics being prescribed to at least 55% of the patients. Among those with wound swab results, Cephalosporin’s, Aminoglycosides and Penicillin’s constituted 85.5% of all antibiotic prescriptions with each being administered to 66.7%, 38.9% and 25% patients respectively. The high prevalence of antibiotic prescriptions given without supporting evidence, coupled with the predominance of three classes of antibiotic among the prescriptions, exhibits potentially inappropriate and/or irrational antibiotic prescription habits in the unit. Inappropriate and irrational prescription of antibiotics are known drivers of AMR due to selection pressure thus these results highlight the clinicians role in development of AMR in the unit.^11,23^

Laboratory diagnostic imprecision was evident from wound swab results of almost 20% of patients (11.8% of wound swab samples) that were identified only by gram stain appearance. Another 34 (55.7%) patients out of the 61 that had a wound swab did not get sensitivity testing done and thus the results were incomplete. Whilst such levels of diagnostic imprecision and incompleteness could partly explain the inappropriate prescriptions by clinicians in our setting, several questions concerning our local laboratory services also arise. Such questions concern the diagnostic capacity of microbiology samples in our local laboratory, level of collaboration between clinical and laboratory personnel and enforcement of quality assurance procedures in the laboratory. These results therefore highlight potential areas of weakness in local laboratory services and potential areas of future research.

Although gram positive bacteria were more prevalent, antibiotic resistance was more common among gram negative bacteria (48.8%) than among gram positive bacteria (26.2%) with Pseudomonas species alone accounting for 48.1% of all drug resistant bacteria. Drug resistant gram negative bacteria also affected a greater proportion of patients (74.1%) when compared to gram positive bacteria which affected 59.2%. The higher prevalence of antibiotic resistance among gram negative bacteria is in accordance with international literature which generally points towards gram negative bacteria as the more important source of AMR.^16,23,24^ It was observed that the prevalence of bacteria resistant to individual antibiotics generally reflected the frequency of use with the highest numbers being among; Aminoglycosides and Aminocyclitols, Penicillin’s and Cephalosporin’s. Although an association between the frequency of systemic antibiotics prescription and development of AMR could not be established due to sparsity of data, the observation is consistent with findings from international studies.^25–27^The observation therefore whilst not conclusive, further highlights the need for rational prescription of antibiotics in order to control the development of AMR.

Although it was not possible to determine where the bacteria and their associated resistance were acquired, the array of commonly isolated bacteria suggests predominance of nosocomial acquisition as opposed to community acquisition of the bacteria.^23,24^ Enterococcus faecium, Staphylococcus aureas, Klebsiella pneumonia, Acinetobacter baumannii, Pseudomonas aeroginosa and Enterobacter (ESKAPE) group of bacteria are the most common multidrug resistant and extensively drug resistant bacteria.^24,28^ These bacteria and their associated drug resistance are primarily nosocomial thus their predominance (63%) among drug resistant bacteria in our study bolsters the case for nosocomial acquisition infections and drug resistance in our setting.^28^ This scenario also further highlights the importance of strengthening antibiotic stewardship and infection control strategies in our setting. Such strategies would be especially important since all drug resistant bacteria identified in this study also appear on the WHO list of priority pathogens.^29^ Identifying the onset of sepsis in burns remains a challenge worldwide due to difficulties in distinguishing burn related and infection inflammatory responses as well as the absence of sensitive and specific methods to distinguish these phenomena in the clinical setting.^13,30–34^ Lack of knowledge on post burn pathological changes among clinicians is probably responsible for the increased antibiotic prescription in the absence of laboratory evidence of bacterial presence and thus contributing towards development of AMR in our setting. Post-burn hypermetabolism and systemic inflammatory response syndrome (SIRS) also result in increased in capillary leakage, hypoprotenaemia, decreased protein binding and augmented renal clearance which results in altered pharmacokinetics and pharmacodynamics of drugs including antibiotics.^25,35,36^ The effects of these changes are more pronounced with hydrophilic drugs(e.g. Beta-lactams, Aminoglycosides, glycopeptides and lipopeptides) which account for 79.6% of antibiotic prescriptions in this study and are often augmented by administration of intravenous fluids or inotropes.^35,36^ Since standard dose calculations are based on relatively healthy populations, these changes are likely to result in suboptimal dosing of antibiotics thus compromising therapeutic outcomes and further contributing towards antibiotic resistance.^35,36^

### Limitations

This study has several weaknesses owing to the retrospective nature of data collection used thus the numeric findings of this study ought to be treated as conservative estimates. From the data collected, it was not possible to distinguish bacterial contamination, colonisation and infection. It was also impossible to distinguish nosocomially acquired bacteria from community acquired bacteria. It was also not possible to establish trends and associations in the distribution of wound bacterial isolated and AMR. In the absence of records pertaining to the methods used to isolate microbes found on the wound surface, it impossible to ascertain if the absence of fungal growth on the wound surface is general reflection of the microbial landscape in the unit or a result of weakness in the materials and methods used in sample collection and processing.

### Recommendations

We would therefore recommend establishment of a multidisciplinary team and enforcement of prescription protocols. The multidisciplinary team should include laboratory technicians and pharmacists in the short term, but also microbiologists in the long term. This will allow holistic patient management, timely processing and communication of good quality laboratory results and active monitoring of trends in antibiotic resistance in the unit. This team can also be utilised to implement antibiotic stewardship and infection management programs as well as training of support staff on ways of minimising interpersonal transmission of infections and AMR. Microbiologists and laboratory technicians that are part of this team would also have an additional role of developing and revising local antibiograms which can be used to guide prescription practices of clinicians. A fully functional multidisciplinary team would ultimately help to achieve optimal utilisation of resources and patient outcomes.

Given the analytical challenges emanating from data sparsity there’s need for conducting robust prospective burns related research projects in order to come up with more accurate estimates on the prevalence of AMR. Current challenges in establishing the presence of sepsis in burns patients in our setting also warrant research into identification of cost effective, sensitive and specific biomarkers that are responsive to therapy and are capable of giving timely results.^37^ Procalcitonin, Pro-B-type natriuretic peptide and hemodynamic changes (systemic vascular resistance index and stroke volume index) have been proposed as potential biomarkers which meet most of the aforementioned criteria.^30,32,35^ To the best of our knowledge, the validity of these biomarkers in an African population is not yet published and there is need to make the biomarkers cost effective for use in low and middle income countries where the incidence of burns and burns-related sepsis is greatest.^3–5^ We postulate that validation and adoption of biomarkers in our setting has the potential of curbing the magnitude of inappropriate antibiotic prescriptions and thus AMR. Early identification of sepsis in patients may also help to alleviate sepsis related morbidity and mortality.

## APPENDICES

**Table 1:** Frequencies of bacteria from wound swab culture.

| ANTIBIOTIC NAME | RESISTANT ISOLATES (N=102) | PERCENTAGE OF RESISTANT ISOLATES |
| --- | --- | --- |
| **AMINOGLYCOSIDES &** |  |  |
| AMINOCYCLITOLS | 24 | 23.5 |
| PENICILLIN'S | 20 | 19.6 |
| CEPHALOSPORINS | 17 | 16.7 |
| CHLORAMPHENICOL | 14 | 13.7 |
| QUINOLONES | 13 | 12.7 |
| SULFONAMIDE | 11 | 10.7 |
| MONOBACTAMS | 10 | 9.8 |
| MACROLIDES | 9 | 8.8 |
| CARBAPENEMS | 4 | 3.9 |
| TETRACYCLINES | 1 | 1.0 |

